# Psychological mechanisms of offset analgesia: The effect of expectancy manipulation

**DOI:** 10.1101/2022.08.09.503102

**Authors:** Tibor M. Szikszay, Wacław M. Adamczyk, Janina Panskus, Lotte Heimes, Carolin David, Philip Gouverneur, Kerstin Luedtke

**Affiliations:** Institute of Health Sciences, Department of Physiotherapy, Pain and Exercise Research Luebeck (P.E.R.L.), Universität zu Lübeck, Luebeck, Germany; Laboratory of Pain Research, Institute of Physiotherapy and Health Sciences, The Jerzy Kukuczka Academy of Physical Education, Katowice, Poland; Institute of Medical Informatics, University of Luebeck, Luebeck, Germany

**Author notes:** These authors contributed equally to this work (Tibor M. Szikszay, Wacław M. Adamczyk).

**Keywords:** offset analgesia, suggestion, endogenous pain modulation, placebo effect, nocebo effect

## Abstract

A frequently used paradigm to quantify endogenous pain modulation is offset analgesia, which is defined as a disproportionate large reduction in pain following a small decrease in a heat stimulus. The aim of this study was to determine whether suggestion influences the magnitude of offset analgesia in healthy participants. A total of 97 participants were randomized into three groups (hypoalgesic group, hyperalgesic group, control group). All participants received four heat stimuli (two constant trials and two offset trials) to the ventral, non-dominant forearm while they were asked to rate their perceived pain using a computerized visual analogue scale. In addition, electrodermal activity was measured during each heat stimulus. Participants in both intervention groups were given a visual and verbal suggestion about the expected pain response in an hypoalgesic and hyperalgesic manner. The control group received no suggestion. In all groups, significant offset analgesia was provoked, indicated by reduced pain ratings (p < 0.001) and enhanced electrodermal activity level (p < 0.01). A significant group difference in the magnitude of offset analgesia was found between the three groups (F_[2,94]_ = 4.81, p < 0.05). Participants in the hyperalgesic group perceived significantly more pain than the hypoalgesic group (p = 0.031) and the control group (p < 0.05). However, the electrodermal activity data did not replicate this trend (p > 0.05). The results of this study indicate that suggestion can be effective to reduce but not increase endogenous pain modulation quantified by offset analgesia in healthy participants.

## Introduction

Endogenous pain modulation has been proposed and is discussed as a leading feature of the nociceptive system that can promote or protect the individual against the transition from acute to chronic pain [1,2]. In general, pain modulation can be assessed through experimental paradigms which are believed to reflect complex inhibitory and faciliatory mechanisms within the neuroaxis [3]. Thus, within the peripheral and central nervous system, a variety of individual transmitters and specific receptor types are involved in the modulation and expression of descending inhibition and facilitation [4]. In the central nervous system, pain can be modulated by cognitive, affective and motivational factors [5] including beliefs and expectations [6]. Furthermore, the efficiency of these modulatory pathways can be assessed in the laboratory setting, using paradigms such as conditioned pain modulation (CPM: the “pain inhibits pain”) [2] and/or offset analgesia (OA).

Offset analgesia can be defined as a disproportionally large pain decrease after a minor noxious stimulus intensity reduction [7]. This test procedure is discussed to indicate the efficiency of the descending inhibitory pain modulation system in humans [8]. For almost 20 years, numerous studies have attempted to identify the processes underlying OA, but the physiological mechanisms have not yet been fully understood. Both peripheral [9–11], spinal [12] and supra-spinal mechanisms [9,13–17] have been shown to contribute to OA. However, few experimental procedures in the past effectively modulated the OA effect. For instance, the modulatory influence of primary afferents [11,18] or secondary noxious stimuli [19] are exceptions rather than common findings expanding our knowledge of OA. In contrast, all pharmacological attempts failed to affect OA [20].

Interestingly, psychological interventions have never been used in the context of OA. It is of clinical interest, whether psychological processes influence endogenous pain modulation, especially since - amongst others-placebo and nocebo manipulations have been shown to effectively decrease [21] or increase [22] pain perception, respectively. For example, it has been demonstrated that by administering a nocebo suggestion prior the application of a noxious stimulus, healthy subjects felt more pain [23]. The putative mechanism of such an intervention relates to expectations [24], which has already been observed in CPM experiments [25,26] but not yet in OA.

This experiment attempted to influence the magnitude of OA by manipulating participants’ expectations using suggestions. In this study, suggestions were used to modulate OA selectively, that is, by changing the pain response in the final temperature phase of the paradigm. Therefore, the á priori hypothesis implied that suggestion would influence the OA effect in a bidirectional manner, i.e., analgesia was expected to be increased or decreased, respectively, compared to a control group that was not exposed to any form of suggestion.

## Materials and Methods

### Study Design

This experimental study was conducted as a randomized controlled trial in which healthy, pain-free participants were randomly divided (counterbalanced) into two intervention groups and one control group. Both intervention groups received either a hypoalgesic or a hyperalgesic suggestion related to the pattern of the subsequently applied heat pain within an OA paradigm. The control group received no suggestion. All participants received the identical information about the exact temperature course of the heat stimuli beforehand. In order to perform the suggestions as authentically as possible, a cover story was told to all participants at the beginning of the study. All participants were blinded to the true purpose of the study. The study was previously approved by the ethics committee of the University of Lübeck (file number: 21-028) and pre-registered in the Open Science Framework (https://osf.io/69eyp). All participants provided oral and written informed consent. An overview of the study design is provided in Fig. 1.

**Figure 1.**
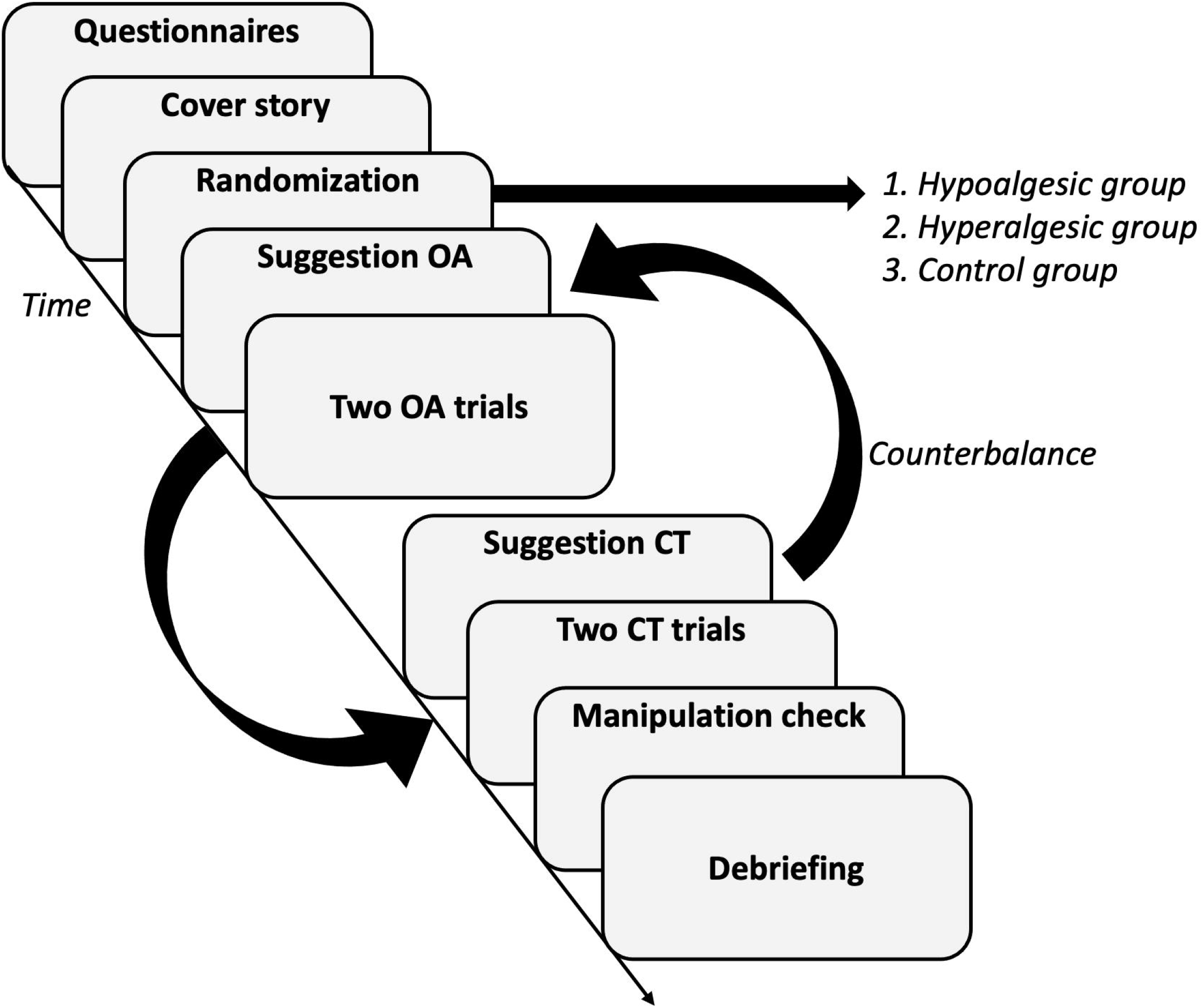
Study design. Before randomization, participants were instructed and a cover story was provided. Participants were told that in this study, changes in electrodermal activity (EDA) would be assessed as a measure of the autonomic nervous system during experimental heat stimuli and used to predict the perception of pain (cover story). Participants were assigned to either i) the hyperalgesic group with verbal suggestion towards no pain analgesia following the temperature reduction (see the example above), ii) the hypoalgesic group with suggestion towards profound analgesia following the temperature reduction or iii) the control group with no intervention. Regardless of the group assignment participants were exposed to two offset and two constant trials provided in a counterbalanced manner.

### Study population

Healthy, pain-free participants aged 18 to 65 years were recruited on the campus of the University of Lübeck. All participants had to confirm that they were healthy and had no cardiovascular, systemic, psychiatric or neurological disease. Furthermore, all participants were excluded if they had a history of chronic pain (> 3 months) within the last 2 years or had experienced pain (including headache, toothache, muscle soreness, etc.) within the last week prior to study participation. In addition, the use of medication, excluding contraceptives, in the last 48 hours was an exclusion criterion. Furthermore, participants were asked not to drink alcohol, exercise, or take pain medication for 24 hours prior to participation in the study and not to drink coffee or smoke cigarettes for 4 hours prior to study participation.

### Equipment

A Pathway CHEPS (Contact Heat -Evoked Potential Stimulator) with a contact area of 27mm diameter was used for the application of the heat stimuli (Medoc, Ramat Yishai, Israel). The thermode was attached to the non-dominant volar forearm approximately 10 cm below the elbow using a blood pressure cuff with a pressure of 25 mmHg. A computerized visual analog scale (COVAS; with the range 0 = “no pain” to 100 = “most tolerable pain”) was used for continuous assessment of pain intensity (Medoc, Ramat Yishai, Israel). Furthermore, during heat stimuli, electrodermal activity (EDA; respiBAN Professional, Plux, Lisbon, Portugal) was measured using two Ag/AgCl hydrogel electrodes (Covidien / Kendall, Dublin, Ireland) at the medial phalanx of index and middle finger of the non-dominant arm (sampling rate of 1000 Hz, PLUX Wireless Biosignals, S.A., Portugal). Electrodermal activity depends on sweat secretion, which is closely related to autonomic nervous system activity [27]. EDA was used to test, if verbal suggestion influences physiological responses and if EDA can be used as an objective marker for OA.

### Experimental heat stimulation

Two constant trials (CT) and two offset trials (OT) were performed on the non-dominant volar forearm, so that the participants were presented with a total of four heat stimuli. The order in which trials were presented was randomized in a counterbalanced fashion. A two-minute pause was kept between each stimulus, during which the thermode was moved on the forearm by a few centimeters. The temperature’s rise and fall rate for all heat stimuli was 15°C/second. During CT, the temperature increased from a baseline level of 35°C to 46°C and remained constant for 40 seconds before returning to the baseline level. During OT, the temperature first increased to 46°C (T1) for 10 seconds, then increased to 47°C (T2) for 10 seconds, and finally decreased again to 46°C (T3) lasting 20 seconds. The temperature pattern of the two trials can be seen in Fig 1. These figures were shown to the participants before the application of the heat stimuli. During the application of the heat stimuli, participants were asked to rate perceived pain continuously and as precisely using the COVAS.

### Suggestion

The hypoalgesic or hyperalgesic groups received suggestions about the expected pain pattern and the pain intensity of the applied heat stimuli. The hypoalgesic group received a placebo suggestion to enhance the effect of OA and adaptation to CT, i.e., to reduce pain perception. The hyperalgesic group, on the other hand, received a nocebo suggestion, which was intended to reduce the effect of OA and adaptation to CT, i.e., to increase pain perception. The expected pain pattern was manipulated verbally (please see S2 Appendix) and supported with the graphical presentation of the assumed pain pattern (Fig S1) and took place directly after the explanation of the temperature gradients, i.e. immediately before application of the respective heat stimulus.

The participants were provided with a cover story. It was explained that the aim of the study was to find out whether the subjective sensation of pain could be “read out” from the physiological reaction of the body (skin conductance) and thus be predicted. This was necessary to also justify the introduction of suggestion as credibly as possible without participants becoming skeptical or biased.

### Questionnaires

Before starting the heat application, participants were asked to complete several questionnaires: The Patient Health Questionnaire (PHQ-9) includes nine questions about depression [28]. While the Pain Vigilance and Awareness Questionnaire (PVAQ) measures pain perception and pain awareness [29], the Pain Sensitivity Questionnaire (PSQ) can be used to determine the subject’s pain sensitivity [30]. The State-Trait Anxiety Inventory-SKD (STAI-SKD) was also collected to determine the participant’s current state anxiety before the experiment [31]. The Social Desirability Scale-17 (SDS-17) was used to measure the participant’s social desirability [32]. Furthermore, the Mindful Attention and Awareness Scale (MAAS) was used to measure dispositional mindfulness [33] and the Life-Orientation Test (LOT-R) was used to measure individual differences between optimism and pessimism based on personality traits [34].

### Manipulation check

To assess the effect of suggestion on pain perception during the OA paradigm, a manipulation check was performed immediately after pain assessment. The following was asked separately for OT and CT: “Please try to recall the moment immediately after receiving heat stimuli. Did you perceive the pain as in the previously displayed figures?” Participants provided a binary (yes vs. no) response.

### Statistical analysis

In the absence of studies investigating OA and verbal suggestion, a meta-analysis examining the effect of verbal suggestion on general pain perception was used to calculate the sample size [22]. With the lowest reported effect size of 0.66 (Cohen’s d), a power of 80%, and an alpha of 0.05, a total number of 30 participants in each group (total = 90) was required (G*Power, University of Düsseldorf [35]) to demonstrate a significant difference between experimental and control groups.

COVAS data from the Medoc software and the EDA signals were synchronized. The time-series data were also sampled to a frequency of 1 Hz. All other statistical analyses were performed using the IBM Statistical Package for Social Science (SPSS version 26, Armonk, NY). The three groups were tested for group differences using age, BMI, sex, dominant hand, and questionnaire data. One-way analysis of variance (ANOVA), Kruskal-Wallis tests, or chi-square tests were used accordingly.

Differences between the groups in their initial pain response at the beginning of the heat stimuli were examined. For this purpose, pain ratings were averaged from the last 5 seconds of the T1 interval. Separately for OTs and CTs (mean value of the two CTs and two OTs) one-way ANOVAs were used to analyze differences between the groups. The primary outcome in this study was the magnitude of the pain response to the T3 interval (OT). To ensure that the magnitude of the OA effect was not under- or overestimated, the mean of 10 seconds centered in the T3 interval (secs. 25 - 34) were extracted. The 5 seconds at the beginning of the interval were not included, because the pain may still decrease during this time, and the 5 seconds at the end of the interval were not included, because the analgesic effect of OA usually decreases after approximately 15 seconds [36]. OT and CT (again mean of the two CTs and OTs) were analyzed separately, as both trials were also separately attempted to be influenced by suggestion. However, dependent t-tests were used to demonstrate whether the pain response and EDA signal from CT were significantly different from OT in each of the groups and thus whether there was an OA effect. To examine the effect of suggestion on OT and CT, a one-way ANOVA was conducted comparing the T3 interval pain response of the three groups as described. The EDA data were analyzed according to identical principles as the pain response. A *p* value of less than 0.05 was considered significant. If statistically significant main or interaction effects were detected, Bonferroni corrected post-hoc t tests were conducted. The correlations between the pain response of the T3 (OT) and the previously described questionnaires were calculated using the Spearman coefficient. No data were missing at the time of analysis.

## Results

A total of 97 participants (hypoalgesic n=32, hyperalgesic n=33, control group n=32) were included in this study. No significant differences were found between groups regarding baseline characteristics (Table 1).

**Table 1:**
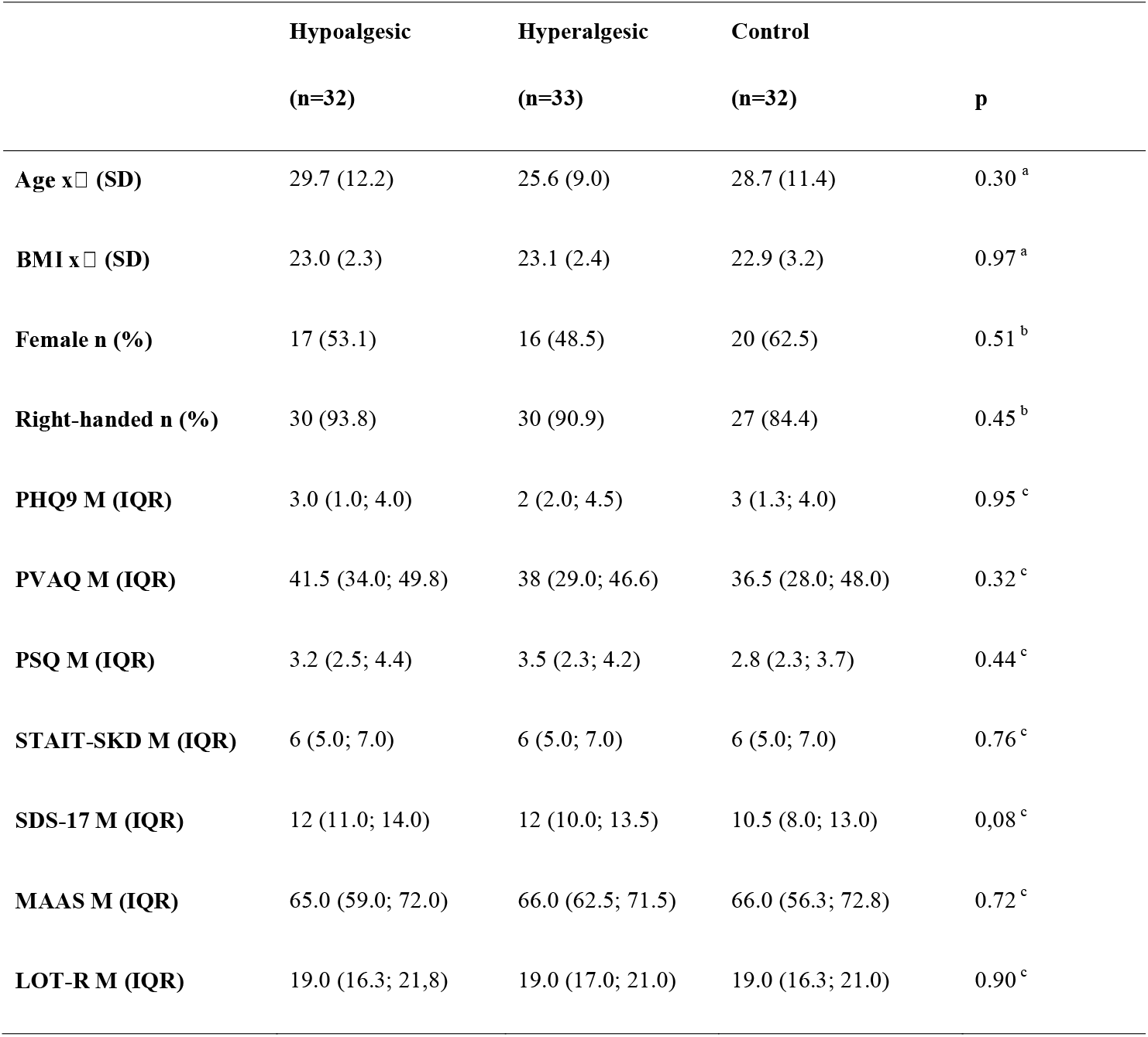
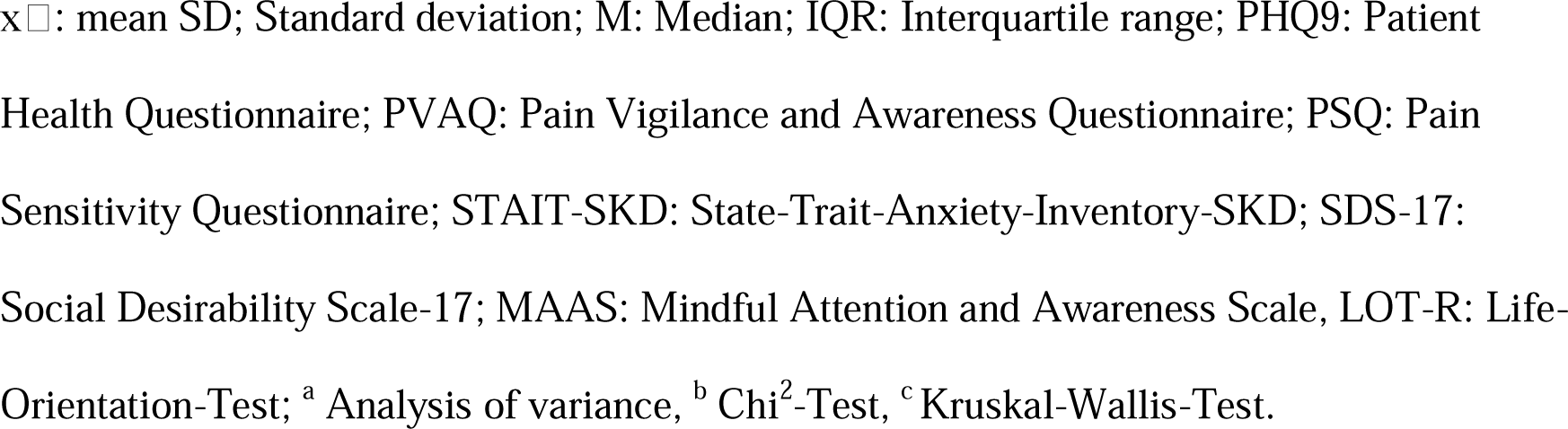
Participant characteristics for each group.

|  | Hypoalgesic | Hyperalgesic | Control |  |
| --- | --- | --- | --- | --- |
|  | (n=32) | (n=33) | (n=32) | p |
| Age x<sup>□</sup> (SD) | 29.7 (12.2) | 25.6 (9.0) | 28.7 (11.4) | 0.30<sup>a</sup> |
| BMI x<sup>□</sup> (SD) | 23.0 (2.3) | 23.1 (2.4) | 22.9 (3.2) | 0.97<sup>a</sup> |
| Female n (%) | 17 (53.1) | 16 (48.5) | 20 (62.5) | 0.51<sup>b</sup> |
| Right-handed n (%) | 30 (93.8) | 30 (90.9) | 27 (84.4) | 0.45<sup>b</sup> |
| PHQ9 M (IQR) | 3.0 (1.0; 4.0) | 2 (2.0; 4.5) | 3 (1.3; 4.0) | 0.95<sup>c</sup> |
| PVAQ M (IQR) | 41.5 (34.0; 49.8) | 38 (29.0; 46.6) | 36.5 (28.0; 48.0) | 0.32<sup>c</sup> |
| PSQ M (IQR) | 3.2 (2.5; 4.4) | 3.5 (2.3; 4.2) | 2.8 (2.3; 3.7) | 0.44<sup>c</sup> |
| STAIT-SKD M (IQR) | 6 (5.0; 7.0) | 6 (5.0; 7.0) | 6 (5.0; 7.0) | 0.76<sup>c</sup> |
| SDS-17 M (IQR) | 12 (11.0; 14.0) | 12 (10.0; 13.5) | 10.5 (8.0; 13.0) | 0.08<sup>c</sup> |
| MAAS M (IQR) | 65.0 (59.0; 72.0) | 66.0 (62.5; 71.5) | 66.0 (56.3; 72.8) | 0.72<sup>c</sup> |
| LOT-R M (IQR) | 19.0 (16.3; 21.8) | 19.0 (17.0; 21.0) | 19.0 (16.3; 21.0) | 0.90<sup>c</sup> |

$\bar{x}$ : mean SD; Standard deviation; M: Median; IQR: Interquartile range; PHQ9: Patient Health Questionnaire; PVAQ: Pain Vigilance and Awareness Questionnaire; PSQ: Pain Sensitivity Questionnaire; STAIT-SKD: State-Trait-Anxiety-Inventory-SKD; SDS-17: Social Desirability Scale-17; MAAS: Mindful Attention and Awareness Scale, LOT-R: Life-Orientation-Test; <sup>a</sup> Analysis of variance, <sup>b</sup> Chi<sup>2</sup>-Test, <sup>c</sup> Kruskal-Wallis-Test.

Mean pain curves and averaged pain from T3 intervals are presented in Figs 2 and 3, respectively. No significant differences were found between all groups regarding the T1 interval for either OT (F_(2, 94)_ = 2.48, p = 0.09, η^2^_p_ = 0.06or CT (F_[2, 94]_ = 1.81, p = 0.17, η^2^_p_ = 0.04). Dependent t-tests showed that OT regarding the T3 interval was significantly different from CT in the pain ratings (hypoalgesic: t_[31]_ = 6.3, p < 0.001, d_z_ = 1.12; hyperalgesic: t_[32]_ = 5.8, p < 0.001, d_z_ = 1.01; control: t_[31]_ = 7.1, p < 0.001, d_z_ = 1.25) as well as EDA (hypoalgesic: t_[31]_ = 4.1, p < 0.001, d_z_ = -0.71; hyperalgesic: t_[32]_ = 4.5, p < 0.001, d_z_ = -0.78; control: t_[31]_ = 3.5, p < 0.01, d=-0.61) in all groups, indicating an OA effect within each of the groups.

**Figure 2.**
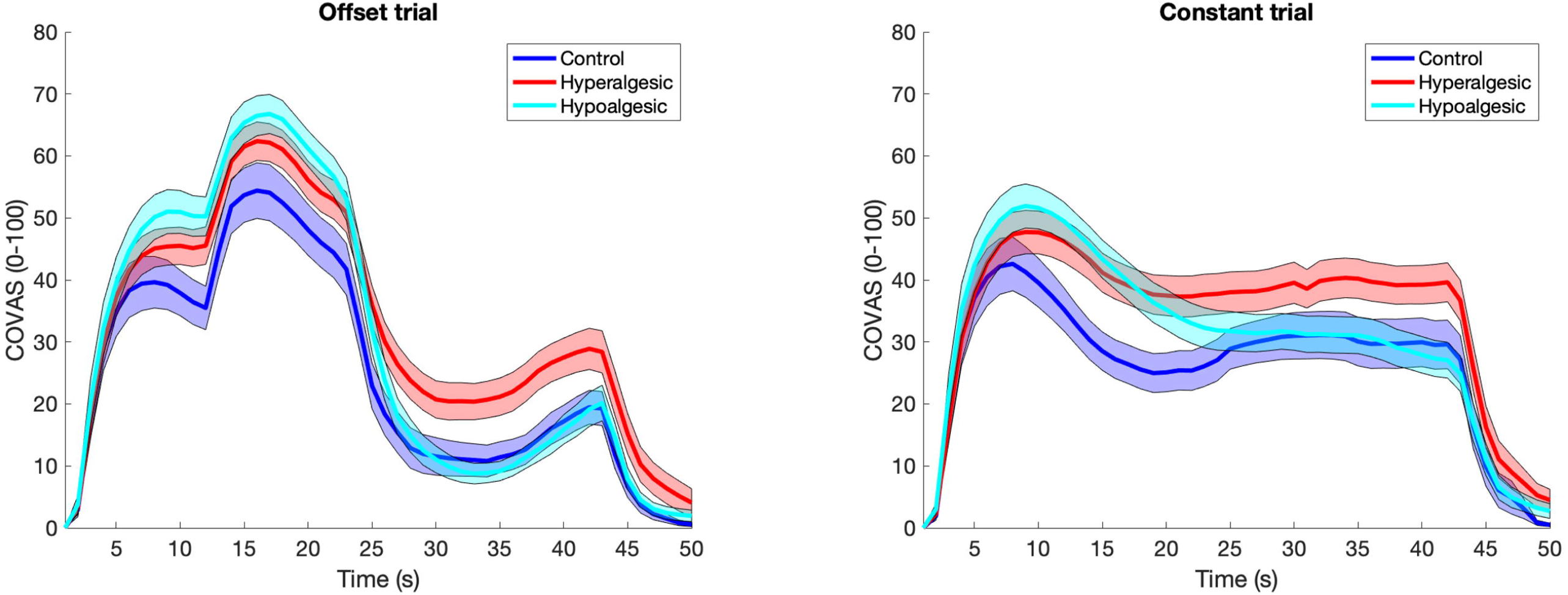
Pain ratings in offset analgesia (left) and constant trials (right). Note that in offset analgesia trials, pain was disproportionally reduced during the last 20s of thermal stimulation assessed via a computerized visual analog scale (COVAS). Hyperalgesic suggestion inhibited the development of profound analgesia present in the control as well as the hypoalgesic group. Suggestion affected constant trials in a similar fashion. Bold curves represent mean pain whereas shaded zones are standard errors of the mean (SEM).

**Figure 3.**
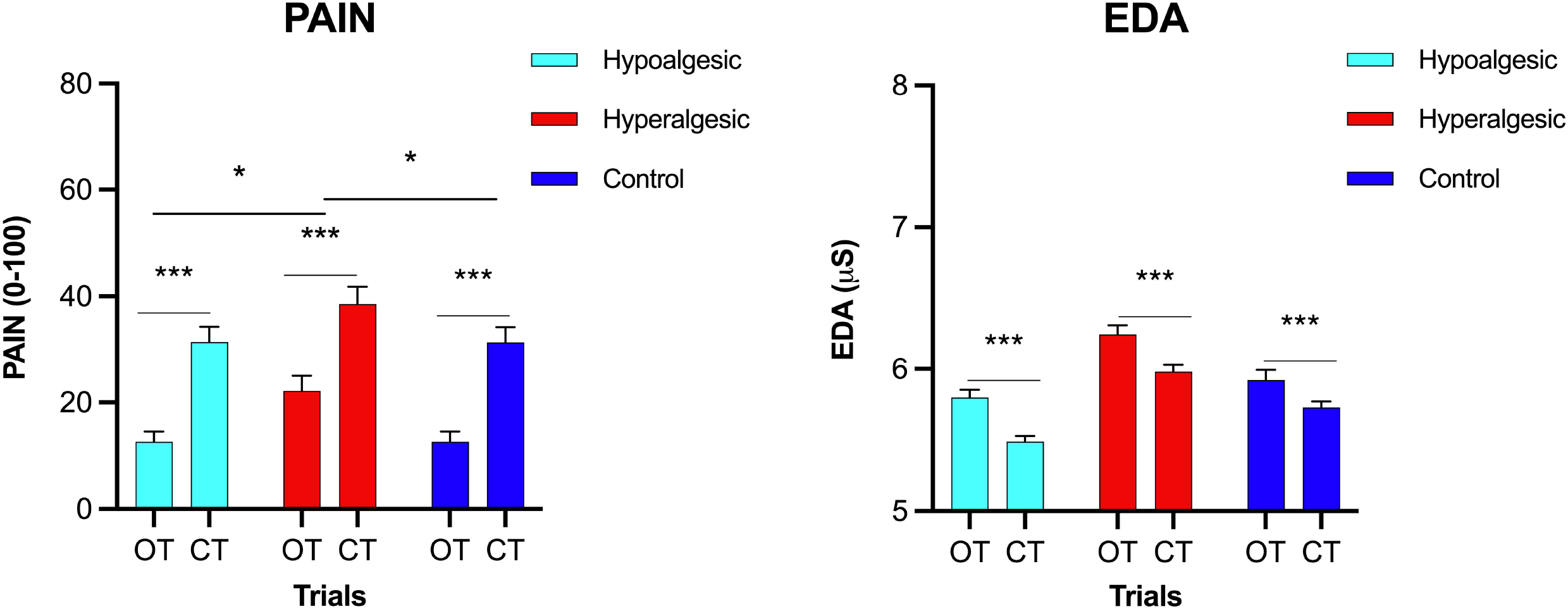
Within- and between-group effects for pain assessed via a computerized visual analog scale (COVAS, left) and electrodermal activity (EDA, right). Offset analgesia was reduced in the hyperalgesic group as reflected by a less pronounced difference in pain (averaged of 25-34s interval) between offset trials (OT) and constant trials (CT). Error bars represent standard errors of the mean (SEM), * indicates a significant difference at p < 0.05, *** p < 0.001

During OT, a significant difference for the factor “group” was found between the three groups in pain ratings at T3 (F_[2, 94]_ = 4.81, p = 0.01, η^2^ _p_ = 0.10). Bonferroni-corrected post-hoc t-tests showed significantly greater pain in the hyperalgesic group than in both the hypoalgesic (p = 0.03) and control (p = 0.02) groups. In contrast, no significant difference was found between the hypoalgesic and the control group (p = 1.00). Regarding the CT (T3 interval), no significant difference was shown between all groups (F_[2, 94]_ = 2.08, p = 0.13, η^2^_p_= 0.13), indicating that the verbal suggestion affected OA trials in the hyperalgesic group and not the pain response to constant trials. Furthermore, no significant difference was shown between the groups regarding EDA in the T3 time interval, neither for OT (F_[2, 94]_ = 0.98, p = 0.38, η^2^_p_ = 0.02) nor for CT (F_[2, 94]_ = 0.91, p = 0.40, η^2^_p_ = 0.02, Figs 3 and 4).

**Figure 4.**
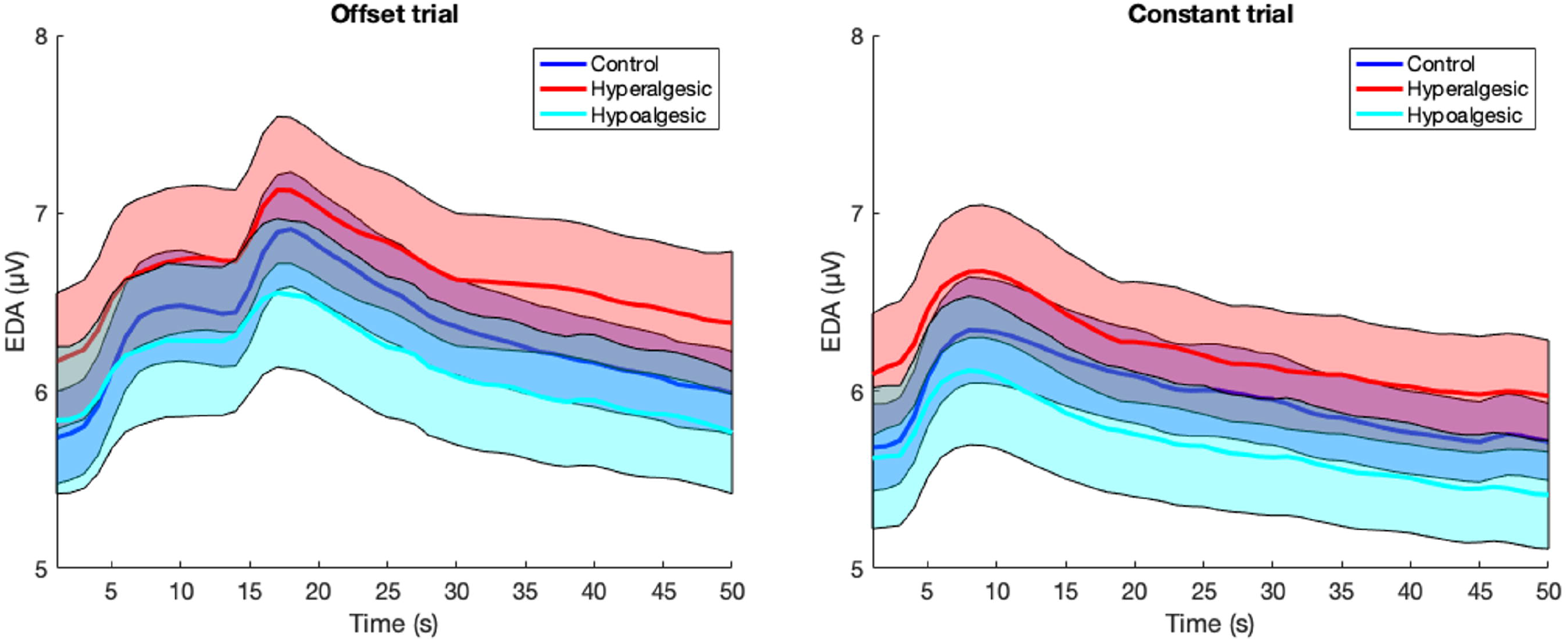
Electrodermal activity (EDA) in offset analgesia (left) and constant trials (right). Compared to constant trials, offset analgesia produced paradoxically higher EDA levels during the T3 interval. Bold curves represent mean pain whereas shaded zones are standard errors of the mean (SEM).

After completion of the study, 70.1% (n = 68) of the participants stated that they perceived pain during OT that was in line with the provided suggestion. This was confirmed by 90.6% (n = 29) of the participants in the hypoalgesic group, but only 21.2% (n = 7) in the hyperalgesic group. A significant difference between groups was observed (𝒳^2^(1, 65) = 31.7, p < 0.001, Φ = 0.70). 74.2% of participants (n = 72) confirmed this for the CT. Thereby, 75.0% (n = 24) confirmed this in the hypoalgesic, but only 48.5% (n = 16) in the hyperalgesic group (𝒳^2^(1, 65) = 4.8, p = 0.03, Φ = 0.27).

No significant correlations were found between the previously described questionnaires and the T3 pain response (OT) in the hypoalgesic group as well as in the control group (p > 0.05, r < 0.3). However, in the hyperalgesic group a significant correlation with the LOT-R was shown (r = -0.45, p < 0.01), which can be attributed mainly to optimism characteristics (optimisms score: r = -0.55, p < 0.01) and not to pessimism characteristics (pessimisms score: r = 0.27 (p = 0.13). All correlation results are presented in the supporting information (S3 Table).

## Discussion

In summary, it can be concluded that OA was provoked in all groups, independent of the verbal manipulation. However, the pain response but not the EDA response during an OA paradigm was influenced by visually reinforced verbal suggestion in healthy participants via nocebo suggestion, but not via placebo suggestion.

### Expectancy mechanism

To the best of our knowledge, this is the first study that has attempted to influence OA using suggestion. However, similar results have already been reported for studies that attempted to influence outcomes using other paradigms to quantify endogenous pain modulation by using suggestion. For example, a similar conclusion was reported by Vaegter et al. (2020) which attempted to influence exercise-induced hypoalgesia (EIH) using suggestion [37]. In that study, EIH was defined as an increased pain threshold and pain tolerance induced by performing a single exercise routine. It was found that volunteers who received a negative suggestion prior to exercise, experienced hyperalgesia instead of EIH. Furthermore, studies found that CPM can also be influenced by suggestions and thereby altered expectation [26,38]. In CPM, the pain response to a painful test stimulus is inhibited by the application of a distant painful conditioning stimulus [39]. Goffaux et al. (2007) studied 20 healthy volunteers regarding their pain perception during the CPM paradigm while they were given different verbal suggestions about the expected pain process [38]. While the placebo group experienced profound analgesia, this was absent in the nocebo group. Moreover, Bjørkedal and Flaten (2012) also found an effect of verbal suggestion on pain perception in the CPM paradigm [26]. It should be noted, however, that OA is not based on the same mechanisms as CPM and EIH, making them not directly comparable. For example, CPM, unlike OA, can be influenced by ketamine [40], and the two paradigms are underlying distinct brain mechanisms [41]. In addition, other studies have shown that there is no correlation between OA and EIH [42] or OA and CPM [41,43], also suggesting individual mechanisms of these pain modulation phenotypes. However, based on these similar results for CPM, EIH, and OA, it is reasonable to assume that altered expectations manipulated by suggestion influence endogenous pain modulation processes as quantified via various paradigms.

In general, it can be assumed that suggestion influences OA, since brain activity during OA overlaps with the activity during placebo analgesia [8,44]. Previous studies have shown that especially the activation of the rostral anterior cingulate cortex (rACC) and the dorsolateral prefrontal cortex (DLPC) play a major role in placebo analgesia [45,46]. They are both functionally connected with the periaqueductal gray and the rostral ventromedial medulla (PAG-RVM system), which can send inhibitory projections to the spine and thereby elicit a diffuse analgesic response [47]. Increased activation of the DLPC and PAG-RVM circuits were also found during OA [13,15,41], suggesting that the mechanisms of placebo analgesia and OA may be similar. However, this is contradicted by the results that placebo suggestion did not produce increased pain reduction in OA in this study. This can be explained by the fact, that previous studies have shown that placebo analgesia is mediated primarily by increased release of endogenous opiates [48], whereas OA has been shown to be opioid-independent [49]. For example, placebo analgesia can be blocked by the opioid antagonist naloxone [50], whereas naloxone, on the other hand, has no effect on the magnitude of OA [49]. One study has found that nocebo hyperalgesia is mediated by the neurotransmitter cholecystokinin (CCK) [51]. However, the effect of CCK on OA has not yet been studied, although it is relevant since the nocebo manipulation could have influenced the OA.

Interestingly, no significant effect of the suggestion was found on CT in this study. Pain perception in the T3 interval of the CT was neither increased nor decreased. This result is in contrasts with other reports because, in principle, both placebo and nocebo effects of verbal suggestion have been found in previous studies to influence a wide variety of noxious stimuli [21,52]. For example, the study by van Laarhoven et al. (2011) found an effect of nocebo verbal suggestion on pain perception in healthy women. However, this study did not use tonic heat stimuli, but mechanical and electrical stimuli [53]. The methodology of other studies also differed in many ways from the present study. For example, studies often used other stimulus modalities (e.g., cold, electric shocks, ischemic pain), or did not use verbal and visual suggestion but used either conditioning alone or a combination of conditioning and suggestion to influence pain perception [21,52]. Furthermore, no study was found that investigated the effect of verbal and visual suggestion on a constant (tonic) heat stimulus, as done in this study. However, one explanation for the differences in influences on CT versus OT could be the difference in physiological processing. It is suggested that pain adaptation to a moderate, constant heat stimulus is primarily mediated by peripheral mechanisms [54–56]. In comparison, both peripheral and central mechanisms are known to shape OA [8]. Since peripheral mechanisms cannot be influenced by suggestion, this could be a possible explanation for the lack of influence on CT. At the same time, the shown suggestibility during OT could support the assumption that OA is primarily a central phenomenon, as pain perception was modulated by expectancy, here. Further studies comparing the suggestibility of responses to OT and CT are needed to draw further conclusions about the underlying mechanisms.

As a limitation, suggestions might not have been fully successful, as shown by the results of the manipulation check. Although the majority (70.1%) of the subjects reported after the completion of the study that they perceived pain during OT according to the given suggestion. However, in contrast to the hypoalgesic group (90.6%), only 21.2% confirmed this in the hyperalgesic group. Thus, it can be assumed that one reason for this may be the exceptionally robust analgesia during the OA paradigm. Thus, these results show that OA can be influenced only unidirectionally, being more likely to be enhanced but more difficult to be inhibited.

### Physiological mechanisms

To the best of our knowledge, this is the first experiment which recorded EDA during OA. This was done for the following reasons: Firstly, OA has been shown to be mediated by the activation of brain areas associated with the regulation of autonomic reactivity [13,15,57,58], thus we aimed to capture this variable continuously to test if OA measurements behaviorally overlap with physiological responses. Secondly, we aimed to investigate if verbal suggestion alters both, subjective and objective outcomes during an OA paradigm.

Interestingly, results of this study showed that, contrary to our prediction, EDA responses to OT were not decreased alongside pain perception. In turn, the T2 temperature increase elevated the EDA level which persisted during the T3 interval, whereas the pain response was reduced. It can be suggested that the higher stimulus in T2 of the OT activates the descending pain inhibition pathways and therefore inhibits pain. In fact, the EDA level during CT gradually decreased over-time which was, in general, associated with higher pain compared to OTs. During OA, endogenous modulatory mechanisms have been shown to be activated [9], which are believed to be driven by PAG activation [13]. PAG is an anatomic structure with multiple nociceptive projections and it plays a crucial role in autonomic control [57,58]. Whether the enhanced activation in PAG explains reversed offset in EDA needs to be determined.

Our psychological manipulation did not influence the EDA signal. Although EDA has been used extensively to capture reported pain intensity and has been shown to be a potent biomarker in pain prediction [59], we could not observe that autonomic reactivity was influenced by verbal suggestion. Similar results have been reported in some of the previous experiments [60], in which verbal suggestions towards analgesia or hyperalgesia were provided [61]. It cannot be, however, excluded that this is a result of the relatively small effect size observed at the behavioral level (pain).

### Psychological mechanisms

The results of this study could serve as an explanatory approach to describe why OA is reduced in chronic pain patients. Various studies showed that a large proportion of chronic pain patients have dysfunctional beliefs about their condition and dysfunctional coping strategies in dealing with their condition [62]. It can be hypothesized that because of these dysfunctional beliefs and coping strategies, chronic pain patients have a fundamentally more negative expectancy toward pain. In this study, a negative expectancy was also evoked in the nocebo group by a suggestion in healthy participants. These participants also subsequently showed reduced OA. Thus, the expectation towards pain could have a decisive influence on the magnitude of OA. For the reduced OA in the nocebo group, the optimism of a person seems to play a role. According to the results, it can be assumed that a more pronounced optimism reduces the effect of suggestion of the nocebo group on OA. Thus, the individual could protect from a more pronounced hyperalgesia (nocebo effect) as a result of the received suggestion. In contrast, no significance of other psychological factors considered for the effect of the suggestions could be observed.

## Supporting information

Visual materials

Verbal suggestions

Correlations

## Acknowledgments

The authors thank the Institute of Medical Informatics, University of Lübeck, kindly for providing the research facilities and equipment.

## Supporting information

**S1 Fig. Schematic representation of the heat stimuli and the suggestion figures of the expected pain perception**. Heat stimuli within the Offset Trial (A): T1 interval (0-9 sec) at 46°C, T2 interval (10-19 sec) at 47°C, T3 interval (20-40 sec) at 46°C. Heat stimuli within the Constant Trial (B): constant at 46°C; suggestion figures of the hypoalgesic group during the Offset Trial (C), pain perception first increases to a level of 50/100, then to 70/100 and drops sharply in the last seconds to an almost non-painful level (approx. 5/100); during the Constant Trial (D), pain perception starts at a level of 50/100 and then slowly and constantly decreases; suggestion images of the hyperalgesic group: during the Offset Trial (E), pain perception first increases to a level of 50/100, then to 70/100 and finally to 50/100 again; during the Constant Trial (F), pain perception remains constant at a level of 50/100.

**S2 Appendix. Standardized verbal suggestions**.

**S3 Table. Correlation analysis of pain scores within the third time interval (T3) and included questionnaires**. PHQ9: Patient Health Questionnaire; PVAQ: Pain Vigilance and Awareness Questionnaire; PSQ: Pain Sensitivity Questionnaire; STAIT-SKD: State-Trait-Anxiety-Inventory-SKD; SDS-17: Social Desirability Scale-17; MAAS: Mindful Attention and Awareness Scale, LOT-R: Life-Orientation-Test, r: spearman-correlation coefficient, p: p-value, significant correlations are marked in bold.

## Notes

### Competing Interest Statement

The authors have declared no competing interest.

## References

1. Ossipov MH, Morimura K, Porreca F. Descending pain modulation and chronification of pain. Curr Opin Support Palliat Care. 2014;8: 143–151. doi:10.1097/SPC.0000000000000055

2. Yarnitsky D. Role of endogenous pain modulation in chronic pain mechanisms and treatment. Pain. 2015;156 Suppl 1: S24–31. doi:10.1097/01.j.pain.0000460343.46847.58

3. Yarnitsky D, Granot M, Granovsky Y. Pain modulation profile and pain therapy: between pro- and antinociception. Pain. 2014;155: 663–665. doi:10.1016/j.pain.2013.11.005

4. Millan MJ. Descending control of pain. Prog Neurobiol. 2002;66: 355–474. doi:10.1016/s0301-0082(02)00009-6

5. Damien J, Colloca L, Bellei-Rodriguez C-É, Marchand S. Pain Modulation: From Conditioned Pain Modulation to Placebo and Nocebo Effects in Experimental and Clinical Pain. Int Rev Neurobiol. 2018;139: 255–296. doi:10.1016/bs.irn.2018.07.024

6. Wei H, Zhou L, Zhang H, Chen J, Lu X, Hu L. The Influence of Expectation on Nondeceptive Placebo and Nocebo Effects. Pain Research and Management. 2018;2018: 1–8. doi:10.1155/2018/8459429

7. Grill JD, Coghill RC. Transient Analgesia Evoked by Noxious Stimulus Offset. Journal of Neurophysiology. 2002;87: 2205–2208. doi:10.1152/jn.00730.2001

8. Hermans L, Calders P, Verschelde E, Bertel E, Meeus M. An Overview of Offset Analgesia and the Comparison with Conditioned Pain Modulation: A Systematic Literature Review. Pain Physician. : 20.

9. Ligato D, Petersen KK, Mørch CD, Arendt-Nielsen L. Offset analgesia: The role of peripheral and central mechanisms. Eur J Pain. 2018;22: 142–149. doi:10.1002/ejp.1110

10. Martucci K, Yelle M, Coghill R. Differential effects of experimental central sensitization on the time-course and magnitude of offset analgesia. Pain. 2012;153: 463–472. doi:10.1016/j.pain.2011.11.010

11. Naugle KM, Cruz-Almeida Y, Fillingim RB, Riley JL. Offset analgesia is reduced in older adults. Pain. 2013;154: 2381–2387. doi:10.1016/j.pain.2013.07.015

12. Sprenger C, Stenmans P, Tinnermann A, Büchel C. Evidence for a spinal involvement in temporal pain contrast enhancement. Neuroimage. 2018;183: 788–799. doi:10.1016/j.neuroimage.2018.09.003

13. Derbyshire SWG, Osborn J. Offset analgesia is mediated by activation in the region of the periaqueductal grey and rostral ventromedial medulla. Neuroimage. 2009;47: 1002–1006. doi:10.1016/j.neuroimage.2009.04.032

14. Nahman-Averbuch H, Dayan L, Sprecher E, Hochberg U, Brill S, Yarnitsky D, et al. Pain Modulation and Autonomic Function: The Effect of Clonidine. Pain Med. 2016;17: 1292–1301. doi:10.1093/pm/pnv102

15. Yelle MD, Oshiro Y, Kraft RA, Coghill RC. Temporal Filtering of Nociceptive Information by Dynamic Activation of Endogenous Pain Modulatory Systems. J Neurosci. 2009;29: 10264–10271. doi:10.1523/JNEUROSCI.4648-08.2009

16. Yelle MD, Rogers JM, Coghill RC. Offset analgesia: a temporal contrast mechanism for nociceptive information. Pain. 2008;134: 174–186. doi:10.1016/j.pain.2007.04.014

17. Zhang S, Li T, Kobinata H, Ikeda E, Ota T, Kurata J. Attenuation of offset analgesia is associated with suppression of descending pain modulatory and reward systems in patients with chronic pain. Mol Pain. 2018;14: 1744806918767512. doi:10.1177/1744806918767512

18. Asplund CL, Kannangath A, Long VJE, Derbyshire SWG. Offset analgesia is reduced on the palm and increases with stimulus duration. Eur J Pain. 2020. doi:10.1002/ejp.1710

19. Szikszay TM, Adamczyk WM, Hoegner A, Woermann N, Luedtke K. The effect of acute-experimental pain models on offset analgesia. Eur J Pain. 2021;25: 1150–1161. doi:10.1002/ejp.1740

20. Larsen DB, Uth XJ, Arendt-Nielsen L, Petersen KK. Modulation of offset analgesia in patients with chronic pain and healthy subjects - a systematic review and meta-analysis. Scand J Pain. 2021. doi:10.1515/sjpain-2021-0137

21. Peerdeman KJ, van Laarhoven AIM, Keij SM, Vase L, Rovers MM, Peters ML, et al. Relieving patients’ pain with expectation interventions: a meta-analysis. Pain. 2016;157: 1179–1191. doi:10.1097/j.pain.0000000000000540

22. Petersen GL, Finnerup NB, Colloca L, Amanzio M, Price DD, Jensen TS, et al. The magnitude of nocebo effects in pain: A meta-analysis. Pain. 2014;155: 1426–1434. doi:10.1016/j.pain.2014.04.016

23. van Laarhoven Aim, Vogelaar ML, Wilder-Smith OH, van Riel PLCM, van de Kerkhof PCM, Kraaimaat FW, et al. Induction of nocebo and placebo effects on itch and pain by verbal suggestions. Pain. 2011;152: 1486–1494. doi:10.1016/j.pain.2011.01.043

24. Colloca L, Miller FG. How placebo responses are formed: a learning perspective. Philos Trans R Soc Lond, B, Biol Sci. 2011;366: 1859–1869. doi:10.1098/rstb.2010.0398

25. France CR, Burns JW, Gupta RK, Buvanendran A, Chont M, Schuster E, et al. Expectancy Effects on Conditioned Pain Modulation Are Not Influenced by Naloxone or Morphine. Ann Behav Med. 2016;50: 497–505. doi:10.1007/s12160-016-9775-y

26. Bjørkedal E, Flaten MA. Expectations of increased and decreased pain explain the effect of conditioned pain modulation in females. J Pain Res. 2012;5: 289–300. doi:10.2147/JPR.S33559

27. Dawson ME, Schell AM, Filion DL. The Electrodermal System. 4th ed. In: Berntson GG, Cacioppo JT, Tassinary LG, editors. Handbook of Psychophysiology. 4th ed. Cambridge: Cambridge University Press; 2016. pp. 217–243. doi:10.1017/9781107415782.010

28. Reich H, Rief W, Brähler E, Mewes R. Cross-cultural validation of the German and Turkish versions of the PHQ-9: an IRT approach. BMC Psychology. 2018;6: 26. doi:10.1186/s40359-018-0238-z

29. Kunz M, Capito ES, Horn-Hofmann C, Baum C, Scheel J, Karmann AJ, et al. Psychometric Properties of the German Version of the Pain Vigilance and Awareness Questionnaire (PVAQ) in Pain-Free Samples and Samples with Acute and Chronic Pain. Int J Behav Med. 2017;24: 260–271. doi:10.1007/s12529-016-9585-4

30. Ruscheweyh R, Marziniak M, Stumpenhorst F, Reinholz J, Knecht S. Pain sensitivity can be assessed by self-rating: Development and validation of the Pain Sensitivity Questionnaire. Pain. 2009;146: 65–74. doi:10.1016/j.pain.2009.06.020

31. Renner K-H, Hock M, Bergner-Köther R, Laux L. Differentiating anxiety and depression: the State-Trait Anxiety-Depression Inventory. Cogn Emot. 2018;32: 1409–1423. doi:10.1080/02699931.2016.1266306

32. Stöber J. The Social Desirability Scale-17 (SDS-17): Convergent validity, discriminant validity, and relationship with age. European Journal of Psychological Assessment. 2001;17: 222–232. doi:10.1027/1015-5759.17.3.222

33. MacKillop J, Anderson EJ. Further Psychometric Validation of the Mindful Attention Awareness Scale (MAAS). J Psychopathol Behav Assess. 2007;29: 289–293. doi:10.1007/s10862-007-9045-1

34. Hinz A, Sander C, Glaesmer H, Brähler E, Zenger M, Hilbert A, et al. Optimism and pessimism in the general population: Psychometric properties of the Life Orientation Test (LOT-R). Int J Clin Health Psychol. 2017;17: 161–170. doi:10.1016/j.ijchp.2017.02.003

35. Faul F, Erdfelder E, Lang A-G, Buchner A. G*Power 3: A flexible statistical power analysis program for the social, behavioral, and biomedical sciences. Behavior Research Methods. 2007;39: 175–191. doi:10.3758/BF03193146

36. Szikszay TM, Adamczyk WM, Luedtke K. The Magnitude of Offset Analgesia as a Measure of Endogenous Pain Modulation in Healthy Participants and Patients With Chronic Pain: A Systematic Review and Meta-Analysis. Clin J Pain. 2019;35: 189– 204. doi:10.1097/AJP.0000000000000657

37. Vaegter HB, Thinggaard P, Madsen CH, Hasenbring M, Thorlund JB. Power of Words: Influence of Preexercise Information on Hypoalgesia after Exercise-Randomized Controlled Trial. Med Sci Sports Exerc. 2020;52: 2373–2379. doi:10.1249/MSS.0000000000002396

38. Goffaux P, Redmond WJ, Rainville P, Marchand S. Descending analgesia--when the spine echoes what the brain expects. Pain. 2007;130: 137–143. doi:10.1016/j.pain.2006.11.011

39. Nir R-R, Yarnitsky D. Conditioned pain modulation. Curr Opin Support Palliat Care. 2015;9: 131–137. doi:10.1097/SPC.0000000000000126

40. Niesters M, Dahan A, Swartjes M, Noppers I, Fillingim RB, Aarts L, et al. Effect of ketamine on endogenous pain modulation in healthy volunteers. Pain. 2011;152: 656– 663. doi:10.1016/j.pain.2010.12.015

41. Nahman-Averbuch H, Martucci KT, Granovsky Y, Weissman-Fogel I, Yarnitsky D, Coghill RC. Distinct brain mechanisms support spatial vs temporal filtering of nociceptive information. Pain. 2014;155: 2491–2501. doi:10.1016/j.pain.2014.07.008

42. Harris S, Sterling M, Farrell SF, Pedler A, Smith AD. The influence of isometric exercise on endogenous pain modulation: comparing exercise-induced hypoalgesia and offset analgesia in young, active adults. Scand J Pain. 2018;18: 513–523. doi:10.1515/sjpain-2017-0177

43. Szikszay TM, Lévénez JLM, von Selle J, Adamczyk WM, Luedtke K. Investigation of Correlations Between Pain Modulation Paradigms. Pain Med. 2021;22: 2028–2036. doi:10.1093/pm/pnab067

44. Koban L, Jepma M, Geuter S, Wager TD. What’s in a word? How instructions, suggestions, and social information change pain and emotion. Neurosci Biobehav Rev. 2017;81: 29–42. doi:10.1016/j.neubiorev.2017.02.014

45. Meissner K, Bingel U, Colloca L, Wager TD, Watson A, Flaten MA. The placebo effect: advances from different methodological approaches. J Neurosci. 2011;31: 16117–16124. doi:10.1523/JNEUROSCI.4099-11.2011

46. Wager TD, Rilling JK, Smith EE, Sokolik A, Casey KL, Davidson RJ, et al. Placebo-induced changes in FMRI in the anticipation and experience of pain. Science. 2004;303: 1162–1167. doi:10.1126/science.1093065

47. Stein N, Sprenger C, Scholz J, Wiech K, Bingel U. White matter integrity of the descending pain modulatory system is associated with interindividual differences in placebo analgesia. Pain. 2012;153: 2210–2217. doi:10.1016/j.pain.2012.07.010

48. Eippert F, Bingel U, Schoell ED, Yacubian J, Klinger R, Lorenz J, et al. Activation of the opioidergic descending pain control system underlies placebo analgesia. Neuron. 2009;63: 533–543. doi:10.1016/j.neuron.2009.07.014

49. Martucci KT, Eisenach JC, Tong C, Coghill RC. Opioid-independent mechanisms supporting offset analgesia and temporal sharpening of nociceptive information. Pain. 2012;153: 1232–1243. doi:10.1016/j.pain.2012.02.035

50. Levine JD, Gordon NC, Fields HL. The mechanism of placebo analgesia. Lancet. 1978;2: 654–657. doi:10.1016/s0140-6736(78)92762-9

51. Benedetti F, Amanzio M, Vighetti S, Asteggiano G. The biochemical and neuroendocrine bases of the hyperalgesic nocebo effect. J Neurosci. 2006;26: 12014– 12022. doi:10.1523/JNEUROSCI.2947-06.2006

52. Petersen GL, Finnerup NB, Colloca L, Amanzio M, Price DD, Jensen TS, et al. The magnitude of nocebo effects in pain: a meta-analysis. Pain. 2014;155: 1426–1434. doi:10.1016/j.pain.2014.04.016

53. van Laarhoven Aim, Vogelaar ML, Wilder-Smith OH, van Riel Plcm, van de Kerkhof Pcm, Kraaimaat FW, et al. Induction of nocebo and placebo effects on itch and pain by verbal suggestions. Pain. 2011;152: 1486–1494. doi:10.1016/j.pain.2011.01.043

54. LaMotte RH, Campbell JN. Comparison of responses of warm and nociceptive C-fiber afferents in monkey with human judgments of thermal pain. J Neurophysiol. 1978;41: 509–528. doi:10.1152/jn.1978.41.2.509

55. Treede RD, Meyer RA, Raja SN, Campbell JN. Evidence for two different heat transduction mechanisms in nociceptive primary afferents innervating monkey skin. J Physiol. 1995;483 (Pt 3): 747–758. doi:10.1113/jphysiol.1995.sp020619

56. Weissman-Fogel I, Dror A, Defrin R. Temporal and spatial aspects of experimental tonic pain: Understanding pain adaptation and intensification. Eur J Pain. 2015;19: 408–418. doi:10.1002/ejp.562

57. Linnman C, Moulton EA, Barmettler G, Becerra L, Borsook D. Neuroimaging of the periaqueductal gray: state of the field. Neuroimage. 2012;60: 505–522. doi:10.1016/j.neuroimage.2011.11.095

58. Makovac E, Venezia A, Hohenschurz-Schmidt D, Dipasquale O, Jackson JB, Medina S, et al. The association between pain-induced autonomic reactivity and descending pain control is mediated by the periaqueductal grey. The Journal of Physiology. 2021;599: 5243–5260. doi:10.1113/JP282013

59. Posada-Quintero HF, Chon KH. Innovations in Electrodermal Activity Data Collection and Signal Processing: A Systematic Review. Sensors (Basel). 2020;20: E479. doi:10.3390/s20020479

60. Geers AL, Fowler SL, Helfer SG, Murray AB. A test of psychological and electrodermal changes immediately after the delivery of 3 analgesic treatment messages. PAIN Reports. 2019;4: e693. doi:10.1097/PR9.0000000000000693

61. Flaten MA, Aasli O, Blumenthal TD. Expectations and placebo responses to caffeine-associated stimuli. Psychopharmacology. 2003;169: 198–204. doi:10.1007/s00213-003-1497-8

62. Howe CQ, Robinson JP, Sullivan MD. Psychiatric and psychological perspectives on chronic pain. Phys Med Rehabil Clin N Am. 2015;26: 283–300. doi:10.1016/j.pmr.2014.12.003

