## Supplementary figures and images for "Psychological mechanisms of offset analgesia: The effect of expectancy manipulation"

### Visual materials

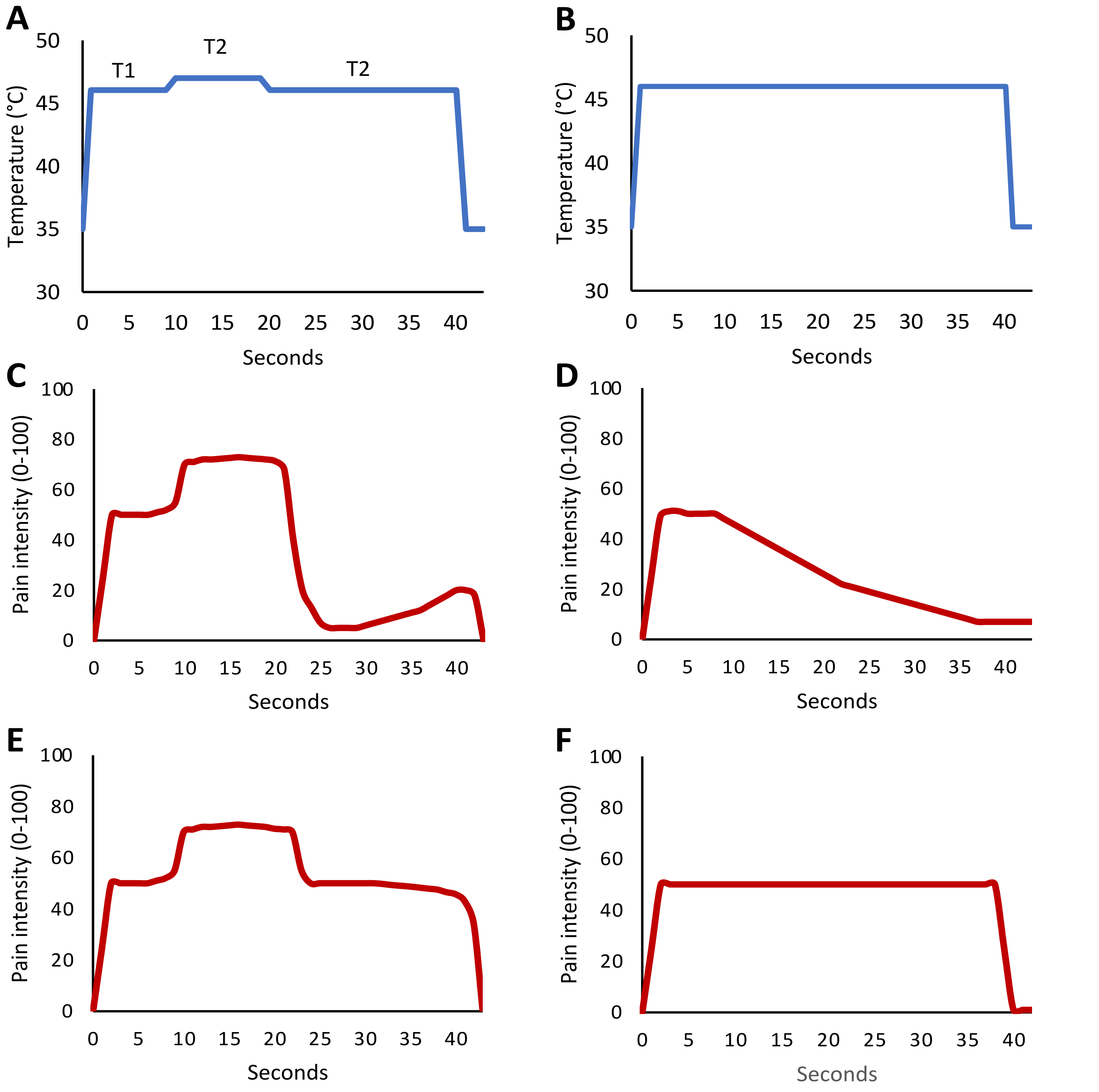
