## Supplementary material for "Psychological mechanisms of offset analgesia: The effect of expectancy manipulation": Verbal suggestions

**S2 - Standardized verbal suggestions**

**Hypoalgesic group**

Offset trial

*"It is an interesting phenomenon that when we receive a heat stimulus that we perceive as painful, we perceive exactly the same heat stimulus with the same temperature following an even stronger heat stimulus as no longer painful at all. This means that the exact same temperature is perceived as pain at first and not during the second time. Several studies have already been conducted with this thermode using exactly these temperature courses. This figure shows the average pain response of the participants to the heat stimulus you are about to receive. Therefore, it's reasonable to expect that you may experience a similar pain perception."*

Constant trial

*"It's an interesting phenomenon that when we receive a heat stimulus that we perceive as painful, over time we become accustomed to that stimulus and gradually perceive it as less painful until eventually it almost doesn't hurt at all. Several studies have already been conducted using this thermode using exactly these temperature trajectories.This figure shows the average pain response from participants to the heat stimulus you're about to receive. Therefore, it's reasonable to expect that you may experience a similar pain perception."*

**Hyperalgesic group**

Offset trial

*"The sensation of pain usually follows temperature pretty closely, as you can see here in the pictures. So, when the temperature increases the pain also becomes more intense, and when the temperature decreases again, logically the pain also drops back to the previous level. Several studies have been done with this thermode using exactly these temperature curves. This figure shows the average pain response of the participants to the heat stimulus you are about to receive. Therefore, it's reasonable to expect that you may experience a similar pain perception."*

Constant trial

*"The sensation of pain usually follows temperature pretty closely, as you can see here in the images. So, if the temperature remains constant as in the following stimulus, then the perceived pain will also remain at a constant level during that. Several studies have been done with this thermode using exactly these temperature patterns. This figure shows the average pain response of the participants to the heat stimulus you are about to receive. Therefore, it's reasonable to expect that you may experience a similar pain perception."*
