## Supplementary material for "Psychological mechanisms of offset analgesia: The effect of expectancy manipulation": Correlations

**S3 Table. Correlation analysis of pain scores within the third time interval (T3) and included questionnaires**

|  | **Hypoalgesic**  **(n=32)** | **Hyperalgesic**  **(n=33)** | **Control**  **(n=32)** |
| --- | --- | --- | --- |
| **PHQ9** |  |  |  |
| r | -0.07 | 0.18 | 0.09 |
| p | 0.7 | 0.32 | 0.62 |
| **PVAQ** |  |  |  |
| r | 0.01 | -0.21 | -0.14 |
| p | 0.95 | 0.24 | 0.44 |
| **PSQ** |  |  |  |
| r | 0.32 | -0.09 | 0.29 |
| p | 0.08 | 0.61 | 0.11 |
| **STAIT-SKD** |  |  |  |
| r | 0.08 | 0.16 | 0.14 |
| p | 0.7 | 0.36 | 0.45 |
| **LOT-R optimism** |  |  |  |
| r | 0.12 | **-0.55** | -0.11 |
| p | 0.52 | **0.001** | 0.57 |
| **LOT-R pessimism** |  |  |  |
| r | -0.04 | 0.27 | -0.10 |
| p | 0.84 | 0.13 | 0.58 |
| **LOTR total** |  |  |  |
| r | 0.08 | **-0.45** | -0.01 |
| p | 0.66 | **0.008** | 0.95 |
| **MAAS** |  |  |  |
| r | 0.01 | -0.28 | -0.30 |
| p | 0.98 | 0.12 | 0.10 |
| **SDS-17** |  |  |  |
| r | -0.10 | -0.19 | -0.23 |
| p | 0.58 | 0.30 | 0.21 |

PHQ9: Patient Health Questionnaire; PVAQ: Pain Vigilance and Awareness Questionnaire; PSQ: Pain Sensitivity Questionnaire; STAIT-SKD: State-Trait-Anxiety-Inventory-SKD; SDS-17: Social Desirability Scale-17; MAAS: Mindful Attention and Awareness Scale. LOT-R: Life-Orientation-Test. r: spearman-correlation coefficient. p: p-value. significant correlations are marked in bold.
